# Precision sampling of discrete sites identified during in-vivo functional testing in the mammalian heart

**DOI:** 10.1101/2023.11.29.569166

**Authors:** Dylan Vermoortele, Camilla Olianti, Matthew Amoni, Francesco Giardini, Stijn De Buck, Chandan K. Nagaraju, Rik Willems, H. Llewellyn Roderick, Karin R. Sipido, Leonardo Sacconi, Piet Claus

**Author notes:** Shared first (Dylan Vermoortele, Camilla Olianti, Matthew Amoni). Shared senior (Karin R. Sipido, Leonardo Sacconi, Piet Claus). Correspondence to Piet Claus, Medical Imaging Research Center, UZ Leuven Herestraat 49, Bus 7006, 3000 Leuven, Belgium. The author deceased while the manuscript was in preparation (Matthew Amoni).

## Abstract

Ventricular arrhythmias after myocardial infarction (MI) originate from discrete areas within the MI border zone (BZ), identified during functional electrophysiology tests. Accurate sampling of arrhythmogenic sites for *ex vivo* study remains challenging, yet is critical to identify their tissue, cellular and molecular signature. In this study, we developed, validated, and applied a targeted sampling methodology based on individualized 3D prints of the human-sized pig heart. To this end, 3D anatomical models of the left ventricle were created from magnetic resonance imaging and fused with biplane fluoroscopy. Regions of interest for sampling were annotated on the anatomical models, from which we created a unique 3D printed cast with custom slits identifying the annotated regions for sampling. The methodology was validated by retrieving ablation lesions created at predefined locations on the anatomical model. We applied the methodology to sample arrhythmia-vulnerable regions after MI during adrenergic stimulation. A novel pipeline of imaging was developed to create a 3D high-resolution map of each sample, highlighting the complex interplay of cellular organization, and altered innervation in the BZ.

---

The power of genetically modifiable mouse models has greatly benefited the understanding of mechanisms and the identification of targets for therapy in cardiovascular disease. Nevertheless, for translation to human, large preclinical models such as the pig or sheep present several advantages, ranging from a similar size to closely related physiology and organ interactions. Owing to their larger scale, these models also allow a more detailed study of regions and structures of interest within the heart. While certain cardiac areas of interest, such as valves or coronary territories can be easily identified morphologically for *ex vivo* studies, it is currently impossible to locate other regions with distinct functional characteristics within the heart after isolation. This is of particular interest for sites of origin of arrhythmias, that can only be identified through functional electrophysiological measurements.

Ventricular tachyarrhythmia is the major cause of sudden cardiac death (SCD) in ischemic HF with reduced ejection fraction, where implantation of an internal cardioverter defibrillator to restore sinus rhythm reduces mortality^1–3^. However, the defibrillator is not completely effective and does not prevent arrhythmias. Unfortunately, pharmacological anti-arrhythmic therapy remains poor with little effective advances over the last 2 decades^4,5^, although pharmacological treatment directed at reducing cardiac remodeling in HF has reduced the incidence of SCD across all etiologies of HF^6^.

A more targeted approach to treating arrhythmias is based on the notion that arrhythmias arise in a limited regional substrate within the heart. Catheter ablation is the destruction of these areas that are responsible for initiating and maintaining the tachyarrhythmias^7^. Whereas initially clinical electrophysiological testing was the only guidance for selecting areas for ablation, this has evolved towards identifying areas that could serve as substrate for arrhythmias^8,9^. In ischemic heart disease, the major substrate for arrhythmias is the region between the infarct scar and the healthy myocardium, the infarct border zone (BZ)^9,10^.

*In vivo* cardiac magnetic resonance imaging (cMRI) of fibrosis and advanced computational analysis of the arrhythmogenic areas in the BZ have led to improved tools to guide ablation^11^. These advances would be strengthened by further insight into the distinctive nature of the arrhythmia-vulnerable areas leading to non-destructive local therapy or target-specific pharmacotherapy.

Additional incentives for a deeper investigation of arrhythmia-vulnerable areas within the BZ emerge from recent work. Clinical and preclinical studies identified a role for altered sympathetic innervation, which is inhomogeneous after MI with areas of altered innervation, that could drive the initiation of arrhythmias as well as their maintenance^12–14^. We recently reported on the heterogeneous electrical remodeling of cardiomyocytes within the BZ, driven by variable gene expression^16^, and we found that premature ventricular complexes elicited by adrenergic stimulation clustered in discrete areas in the BZ^10,15^. While it is feasible to identify and excise the BZ in an isolated heart, a technique to discretely recover *in vivo* identified arrhythmogenic sites within the BZ for mechanistic *ex vivo* characterization is not available.

Hereto, we develop a targeted sampling method to recover *in vivo* identified regions of interest within the pig heart and use it to study discrete arrhythmogenic sites identified after a functional study *in vivo*. Further, we explore the feasibility of detecting structural differences that may occur during the remodeling process in arrhythmogenic versus non-arrhythmogenic sites. We build on our previously developed augmented reality fluoroscopy system (Leuven Augmented Reality Catheter Ablation, LARCA) combined with cMRI^17,18^ by adding a mapping system to tag endocardial sites of interest. The sampling of tagged sites is then enabled by constructing an individualized 3D-printed cast. Given the importance of the autonomic nervous system in the pathophysiology of cardiac arrhythmias and as possible target of therapy^19^, we develop a pipeline to reconstruct in 3D at high-resolution both collagen organization, and the morphometry of adrenergic innervation in BZ samples, building on recent advances of mesoscale tissue imaging^20–22^.

## RESULTS

### Individualized 3D print of the heart for targeted sampling based on *in vivo* imaging

We developed a pipeline for tissue sampling of *in vivo* identified regions of interest. It was validated by determining the accuracy of recovery of small RF lesions at predefined locations in the LV of healthy pigs (Supplemental Figure 1A). First, an *in vivo* baseline cMRI was obtained. From these images, we derived a 3D endocardial model (Figure 1A and Supplemental Figure 2) and defined 9 positions as target sites for sampling distributed in the anterior, posterior and septal wall (Figure 1B). This model provided all the anatomical information needed to construct an individualized 3D printed cast of the heart that included slits for sampling. The epicardial whole heart (left and right ventricle) cast was obtained by outwardly extruding the epicardial cMRI-based model (Figure 1C left and Supplemental Figure S3). The extruded model was divided in two pieces (left and right) cut longitudinally at the level of the interventricular groove. To allow sampling from septal sites after resecting the right ventricle an additional septal plate was derived from the segmented right-sided septum. To avoid misalignment, it was mounted on a plate that exactly fitted the inside of the left epicardial cast (Supplemental Figure S3). Since the cast was based on the original cMRI the pre-defined coordinates on the endocardial surface model were used to create slits for future targeted sampling. Specifically, a beam of 1×1 cm cross-section was centered at each sampling site and oriented perpendicular to the endocardial surface (Figure 1C middle). Targeted sampling slits were created by performing a Boolean operator between the cast and the beams at the sampling sites (Figure 1C right).

**Figure 1.**
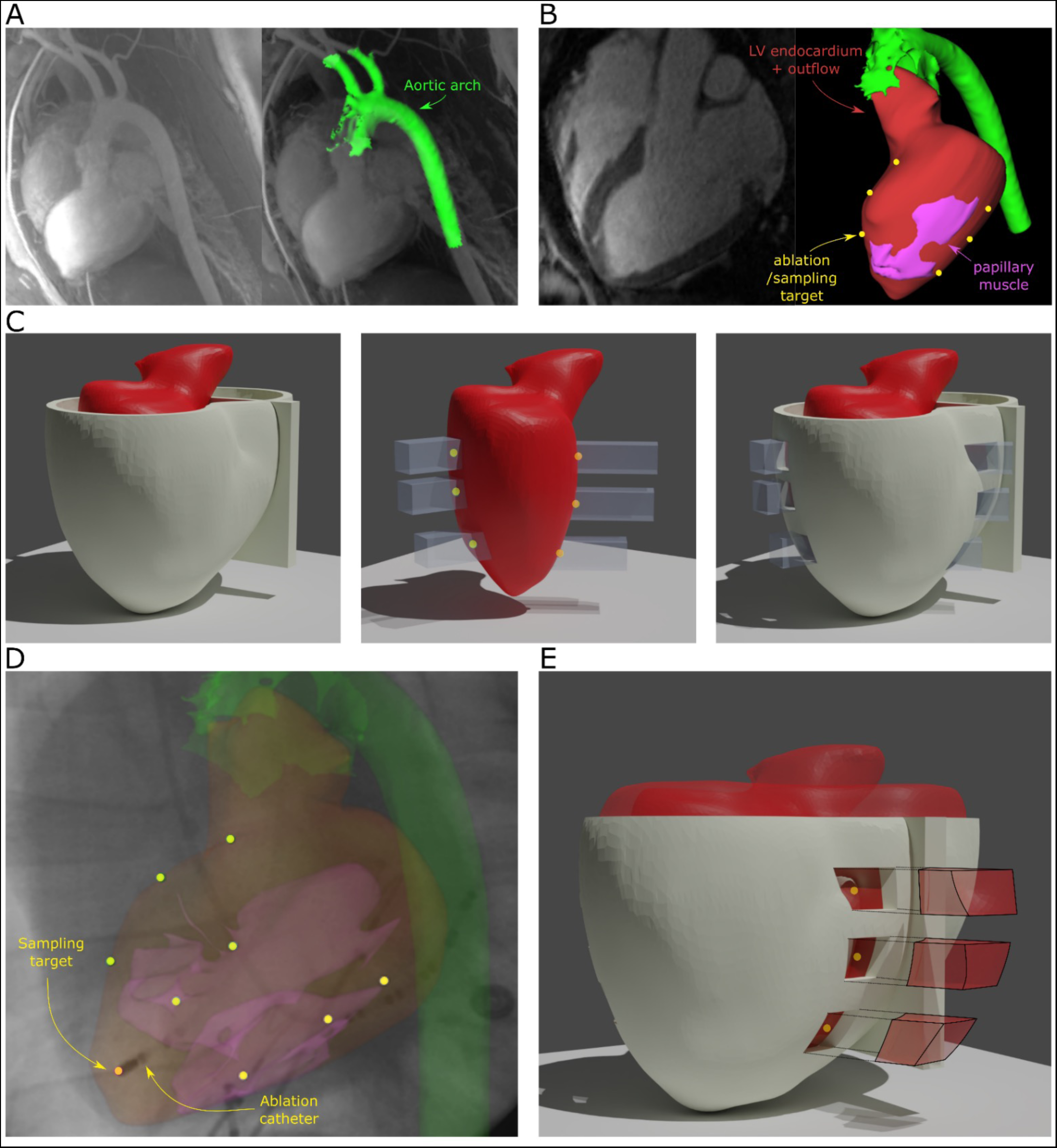
Validation of targeted sampling methodology using cardiac ablation lesions. **A+B)** Pre-operative cardiac magnetic resonance imaging (cMRI) was used to construct anatomical models of the aortic arch and the left ventricle. Positions for ablation lesion creation, and subsequent sampling, were defined on the 3D anatomical model (yellow markers). **C)** An individualized 3D printed cast was constructed using the epicardial segmentation of the *in vivo* cMRI. Slits for targeted sampling were inserted in the model to facilitating the *ex vivo* marking of sampling locations. **D)** Ablation lesions were created under augmented fluoroscopic guidance at the target locations. **E)** The targeted sampling methodology was validated by assessing the accuracy of ablation lesion retrieval.

### Augmented bi-plane fluoroscopy for accurate localization of target sites

In a next step, pigs underwent a percutaneous left ventricular catheterization and we marked the target sites *in vivo* for sampling, by inducing a radiofrequency (RF) lesion (6 mm diameter) centred at the previously defined positions at the anterior, septal and lateral wall. To guide the catheter to these positions, we used the augmented bi-plane fluoroscopy, to fuse the 3D cMRI-derived model and the marked target sites, with the fluoroscopic image (Figure 1D). To validate precision of the catheter guidance, we confirmed the accuracy of aligning the cMRI model with the bi-plane fluoroscopy using edge matching technique (Supplemental Figure 4). Additionally, the augmented bi-plane fluoroscopy simplified catheter navigation by providing anatomical landmarks. In 24 of the pre-defined positions (3 predefined sites could not be reached by the ablation catheter), a small RF ablation lesion of 6 mm diameter was created. At the end of the procedure, the heart was removed and placed in its individualized 3D cast (Figure 1D).

### Assessment of accuracy of 3D print-guided *ex vivo* sampling

The individualized design minimized the freedom to position the refilled heart inside the 3D printed cast and the provided slits explicitly define where to mark the endocardial anterior and lateral sites for sampling (Figure 2A). Removing the right ventricle allowed to mark the RV septal sampling sites using the septal plate. After all the sites were marked, the LV was removed from the cast (Figure 2B). The cavity was opened and placed endocardial surface down to excise the marked sampling sites (Figure 2C). The samples and lesions were measured and photographed and quantified using ImageJ (NIH, USA). Figure 2D-E shows the accuracy and retrieval of the *in vivo* defined sites. For targets of ~ 6mm diameter, using a sample size of 0.88 ± 0.18cm^2^ (Figure 2F) we achieved an accuracy of 2.1 ± 1.7 mm when measuring the center of the target lesion to the center of the sample. This resulted in 89 ± 19.8% of the target being recovered within the sample (Figure 2G).

**Figure 2.**
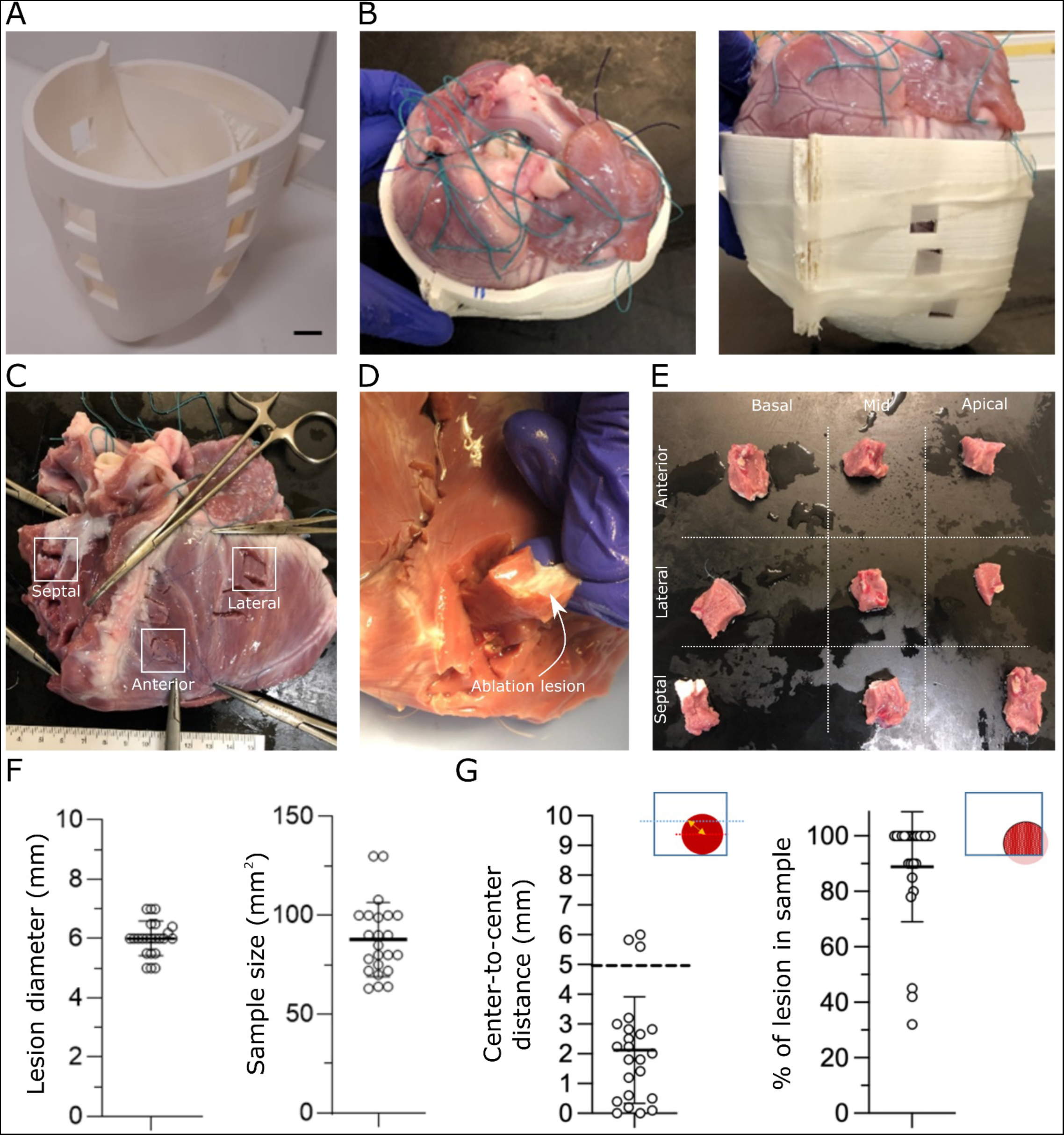
Accuracy of cardiac ablation lesion retrieval using individualized 3D printing based targeted sampling. **A)** An individualized cast model with slits for targeted sampling was 3D printed. **B)** The ex-vivo heart was refilled and fitted inside of the 3D printed cast allowing to mark targets for sampling. **C+D)** The cavity was opened and placed endocardial surface down and the marked sampling sites were excised. **E)** The samples were photographed and the accuracy was quantified. **F)** Ablation lesion size and sample size are given. **G)** The accuracy and percentage of ablation lesion retrieval is illustrated.

### Localization of high frequency PVC sites

The method was applied in a pig model of myocardial infarction to recover arrhythmic sites identified as sites of PVC clustering during an incremental adrenergic stimulation (Supplemental Figure 1B). For this, 4 weeks after creating the MI, a late gadolinium enhanced cMRI was performed, and used to create a 3D model off-line that included the infarct scar (Figure 3A). Two days later the animals underwent an electrophysiology study with augmented bi-plane fluoroscopy, aligning the 3D model with the fluoroscopic images (Figure 3B) An electro-anatomical map was constructed (Figure 3C) and animals received incremental doses of isoproterenol, during a continuous non-contact EP recording. After recording, the origins of PVCs were annotated on the electro-anatomical map (Figure 3D). The majority of PVCs originated from discrete areas within the BZ, consistent with previous findings (Figure 3E)^10^. Subsequently, clusters of frequent PVCs were annotated on the 3D anatomical model using the live augmented bi-plane fluoroscopy by guiding the catheter to the identified PVC site on the electro-anatomical map (Figure 3F). Additionally, a BZ region where no PVCs originated (NO-PVC site) was annotated, as well as a region in the remote, non-infarcted myocardium (remote site).

**Figure 3.**
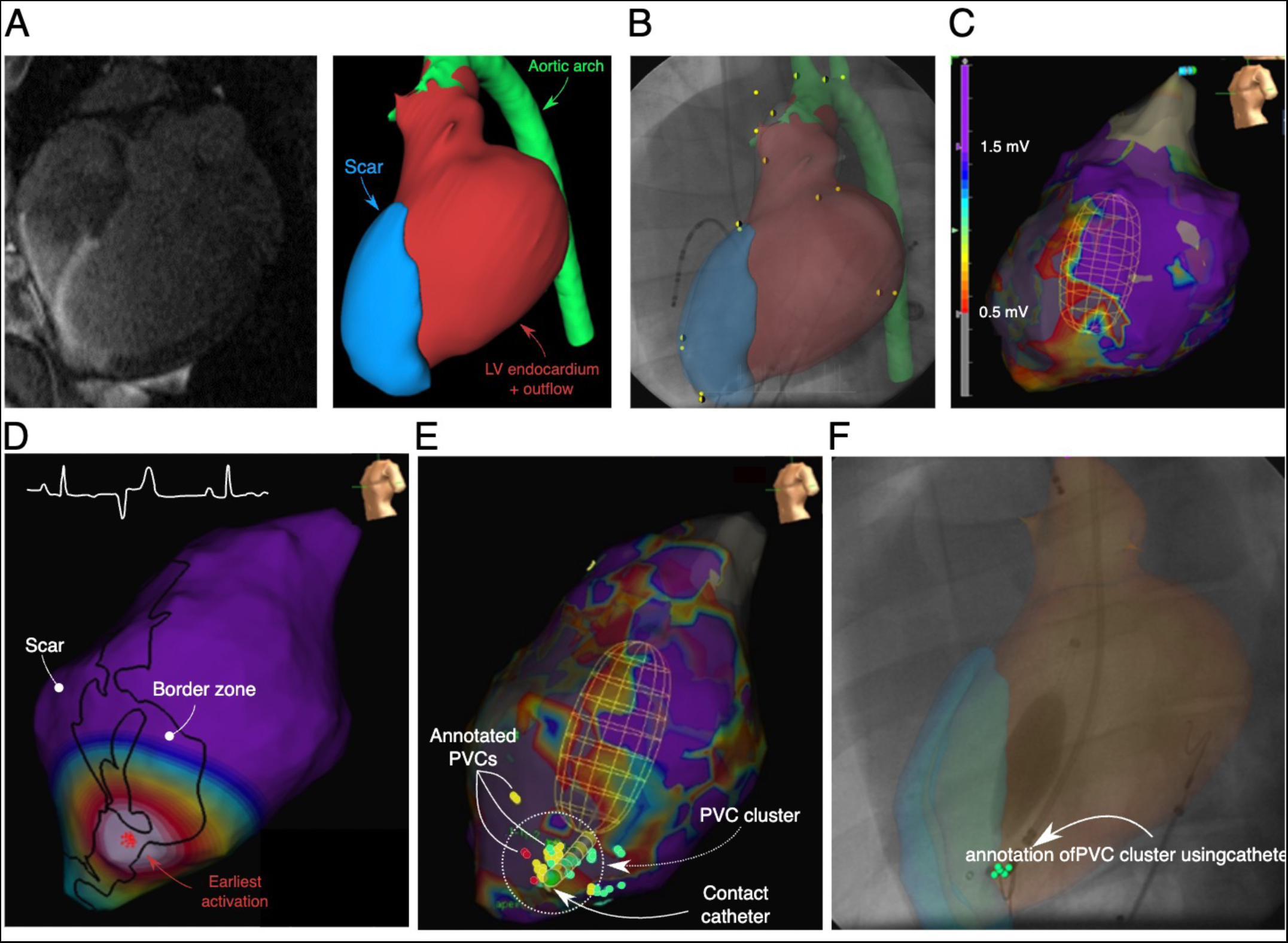
In vivo identification and annotation of arrhythmia vulnerable substrate using augmented bi-plane fluoroscopy and electro-anatomical mapping. **A**) Anatomical models of the left ventricle endocardium, the myocardial infarction and the aortic arch were constructed based on in-vivo magnetic resonance imaging. **B**) The anatomical models were co-registered with bi-plane fluoroscopic imaging with iterative closest point (yellow/green dots) cloud registration **C**) A contact mapping model of the left ventricle is constructed. The bipolar peak-to-peak voltage is used to delineate the infarct (<0.5 mV) and border zone (between 0.5 mV and <1.5mV) regions on the contact model. **D**) The origin of premature ventricular complexes (PVCs) were annotated during incremental adrenergic stimulation using non-contact mapping. **E**) Cluster of PVCs were identified during incremental adrenergic stimulation **F**) A contact mapping catheter was guided to the PVC cluster site and was annotated on the anatomical model using the augmented bi-plane fluoroscopy.

After the electrophysiology study, a 2-day recovery allowed a complete washout of anesthetics and isoproterenol and time to print the 3D individualized cast with sampling slits identifying PVC, NO-PVC and remote regions.

### *Ex vivo* sampling of arrhythmia sites in the MI border zone

Figure 4A shows examples of casts for MI animals, which had sites with high PVC intensity within the BZ, with slits for sampling of remote, PVC and NO-PVC regions of interest, as prepared before the sacrifice. After harvesting of the heart, the ventricles were refilled to the volume matching the *in vivo* cMRI and high-resolution *ex vivo* gadolinium enhanced MRI was performed to assess the anatomy of the heart positioned inside the cast and its correspondence with the in vivo conditions. As a first validation, these ex-vivo images confirmed that all BZ sampling slits (N = 12 in 6 animals) were positioned at the transition from infarct to surviving tissue and remote sites in healthy tissue (N=6 in 6 animals) (Figure 4B). This qualitatively reflects that individualized 3D prints had a high accuracy for identifying sampling regions. Furthermore, we quantified the cardiac filled volumes while the heart was inside the 3D printed cast (Figure 4C). The obtained filling ratio between *in vivo* and *ex vivo* LV volume was 103 ± 25% (Figure 4C), indicating that *in vivo* volumes can be replicated *ex vivo*. Furthermore, the epicardial volume of the heart filled 95.0 ± 2.6% of the available cast volume. The individualized cast thus almost perfectly coincides with the epicardium of the filled heart.

**Figure 4.**
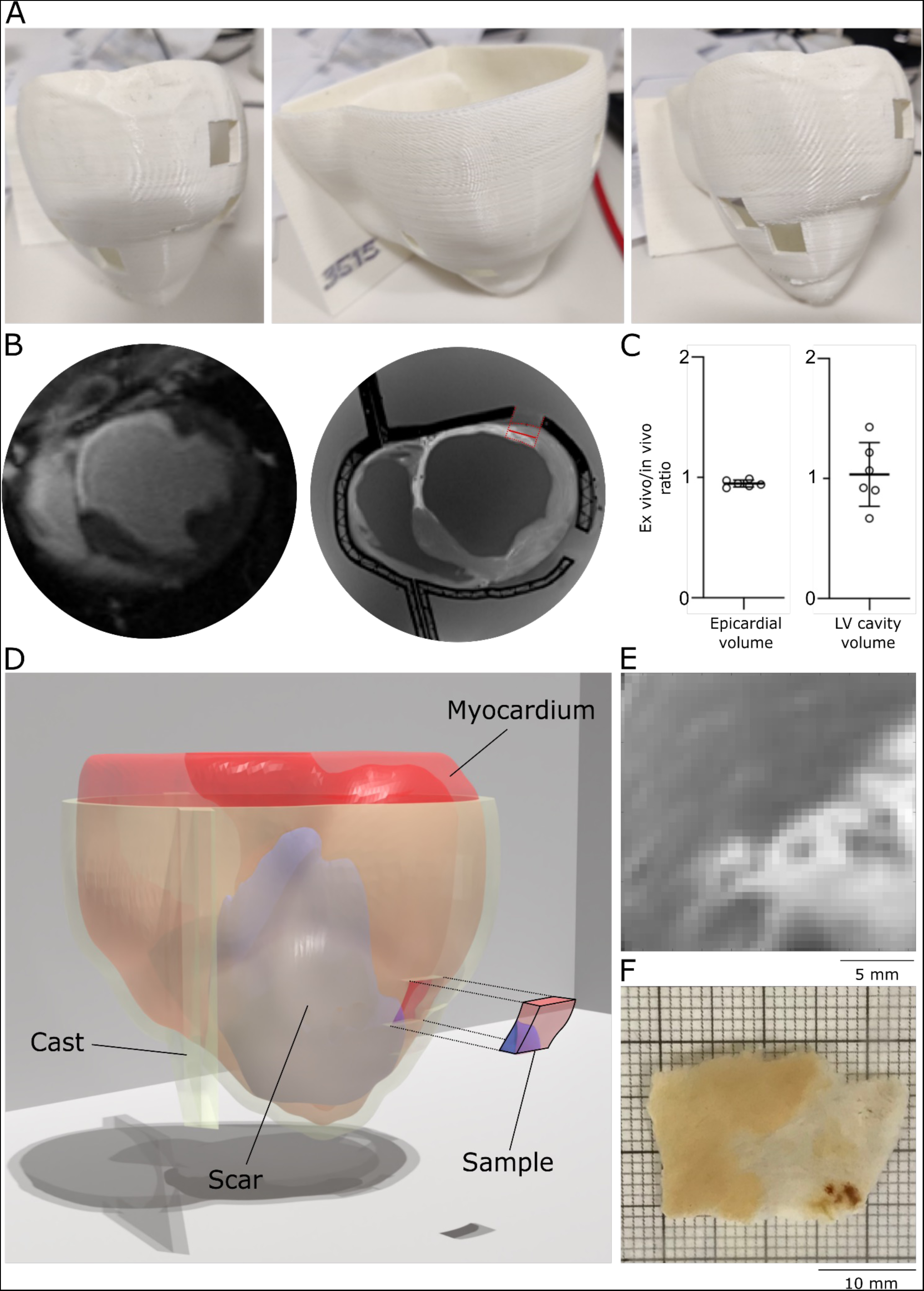
Sampling of in vivo identified arrhythmia vulnerable sites. **A)** A cast model with slits targeting in-vivo identified arrhythmia vulnerable sites was 3D printed. **B)** High resolution *ex vivo* MRI (right) was performed and compared with *in vivo* MRI. **C)** The epicardial volume and left ventricle cavity volume were compared based on *in vivo* and *ex vivo* MRI. **D)** The regions of interest were sampled. **E+F)** The scar structure of the sample was visually compared with the corresponding region on gadolinium enhanced *ex vivo* MRI.

Samples were excised from the MI heart as shown in Figure 1E, with samples extended 4-5 mm in the circumferential direction. To further assess the precision of recovery, we compared the fixed sample, where fibrotic tissue in the infarct is paler than viable myocardium (Figure 4E), with the corresponding image of the ex vivo MRI with LGE image of the infarct in white (Figure 4F). These areas are highly similar, validating that the process of marking and excising the sample does not induce any considerable deviation (Supplemental Video V1).

Samples from three out of the six animals with the largest MI and most frequent PVCs, were selected for developing a pipeline for in depth structural analysis of tissue composition and innervation. From each of these animals a sample from a site in the BZ with clustering of PVCs (BZ PVC) and a sample from a site in the BZ without PVCs (BZ NO-PVC) were processed. In addition, two control samples isolated from the myocardium distal from the infarction were isolated from the first two hearts and were processed as reference (REMOTE). Of these samples, we cut 500 µm thick slices of which we used the first subendocardial slice for analysis.

### Large-area 3D imaging of CLARITY-cleared pig heart slices

3D micron-scale resolution imaging of the slices of (1 × 2 cm) was performed employing non-linear microscopy in combination with a CLARITY-based tissue clearing protocol. CLARITY-related out-of-plane distortions of the specimens were restrained by employing an incubator chamber to maintain the planeness of the slices during the clearing procedure (Figure 5A). The planar expansion that the tissue underwent during the clarification (Figure 5B) was characterized by measuring the expansion coefficients across longitudinal and transversal axes in both myocardial and scar regions (Figure S5). The entire 3D volume of the slices was reconstructed at high resolution using a non-linear microscope by acquiring separate tiles, subsequently stitched together (Figure 5C): cellular membranes are displayed in red (Wheat-Germ Agglutinin, WGA), sympathetic innervation network in green (anti-tyrosine hydroxylase, TOH) and collagen in blue (second-harmonic generation). We conclude that our imaging pipeline allows reliable structural reconstructions of the complete innervation network at single-neuron resolution across entire centimeters-size slices, in both remote and infarcted areas.

**Figure 5:**
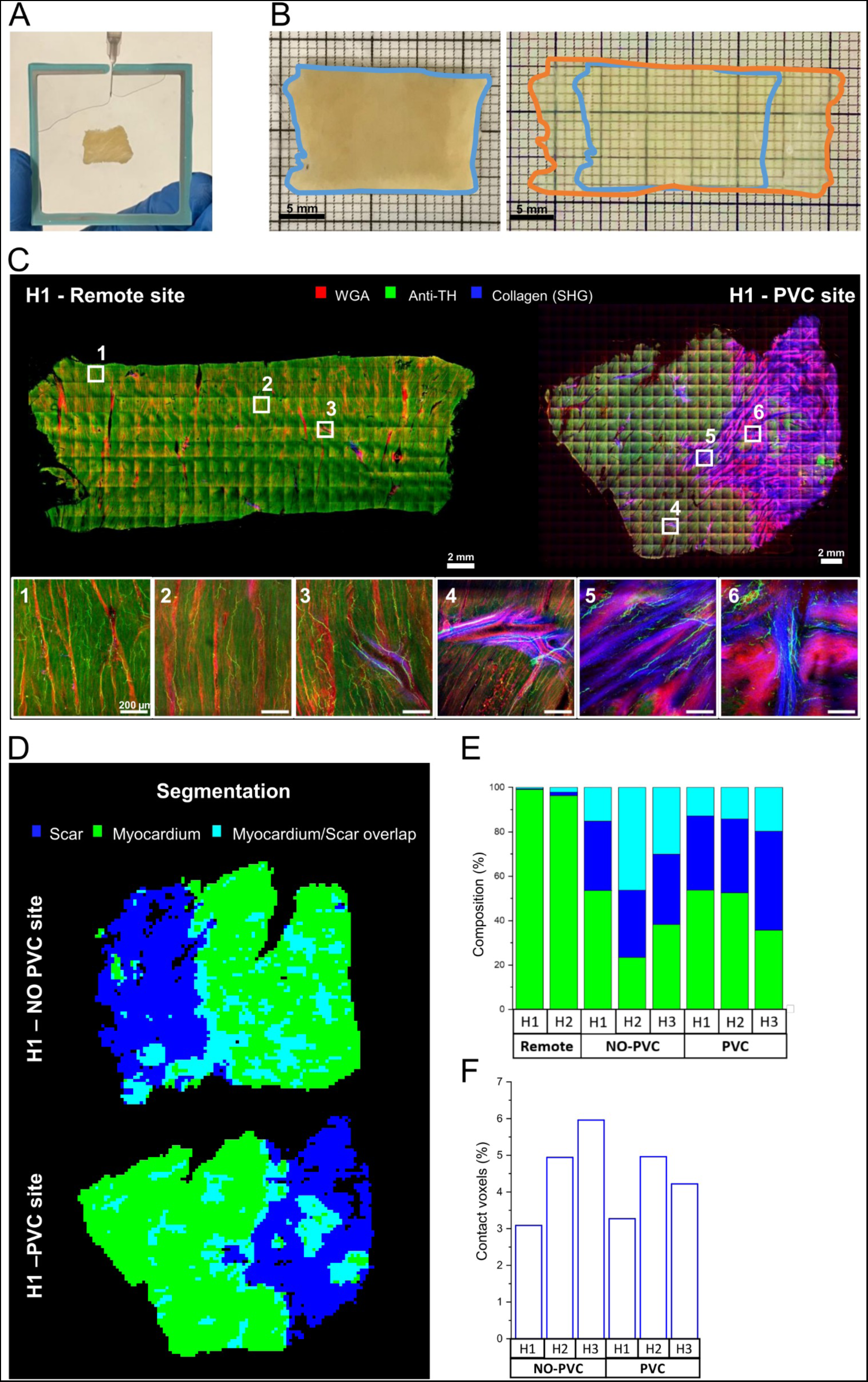
Large-area 3D imaging of CLARITY-cleared pig heart slices. **A)** The custom-made chamber device developed to prevent out-of-plane distortions of the samples during CLARITY-base tissue transformation. **B)** A remote site section before (in PFA) and after CLARITY (in EasyIndex). Borders, before and after clearing, are highlighted in blue and orange respectively. **C)** Maximum intensity projections (MIPs) of 50 (m of a remote and a PVC site (H1): Wheat-Germ Agglutinin (WGA) in red, Anti-tyrosine Hydroxylase (TH) in green and collagen (by Second-Harmonic Generation; SHG) in blue. Three representative tiles for each of the two mosaics are shown in the insets below at full resolution. **D)** Segmented masks of the border-zone samples (H1) composition: myocardial, scar and overlap regions are shown respectively in light green, dark blue and light blue. **E)** The volumetric percentages of the myocardial, scar and overlap regions of all the reconstructed samples. **F)** Percentage of collagen/myocardium contact voxels over the total voxels in all the border-zone samples. Values expressed in 10^-7^.

Differences in collagen distribution are clearly visible between the slices sampled from remote and infarcted zones. The percentage of the myocardial tissue (green), scar tissue (blue) and overlapping region (cyan) over the total volume were quantified by correcting the reconstructed volumes by the estimated expansion coefficients (Figure 5D-E). The data highlight the signature collagen content in the fibrotic patches in the BZ samples, but without clear differences in the PVC vs NO-PVC samples. The percentage of scar-myocardium contact surface area over the total volume of the samples was calculated with no variations between NO-PVC and PVC sites detected (Figure 5F)

### Sympathetic innervation morphometry

Morphometry of the sympathetic innervation network was investigated across slices in a semi-3D fashion by a novel automated software for the detection of nerve endings at full resolution. The core of the analysis is a machine-learning-based discrimination of neurons over the wide variability of signal-to-noise ratio across slices (Figure 6A). The performances of the developed software in removing the background noise and enhancing the signal coming from the nerve endings are represented in Figure 6B. The processed images were then used to assess the local neuronal density and co-alignment of the nerve endings and map them across remote and BZ slices (Figure 7).

**Figure 6:**
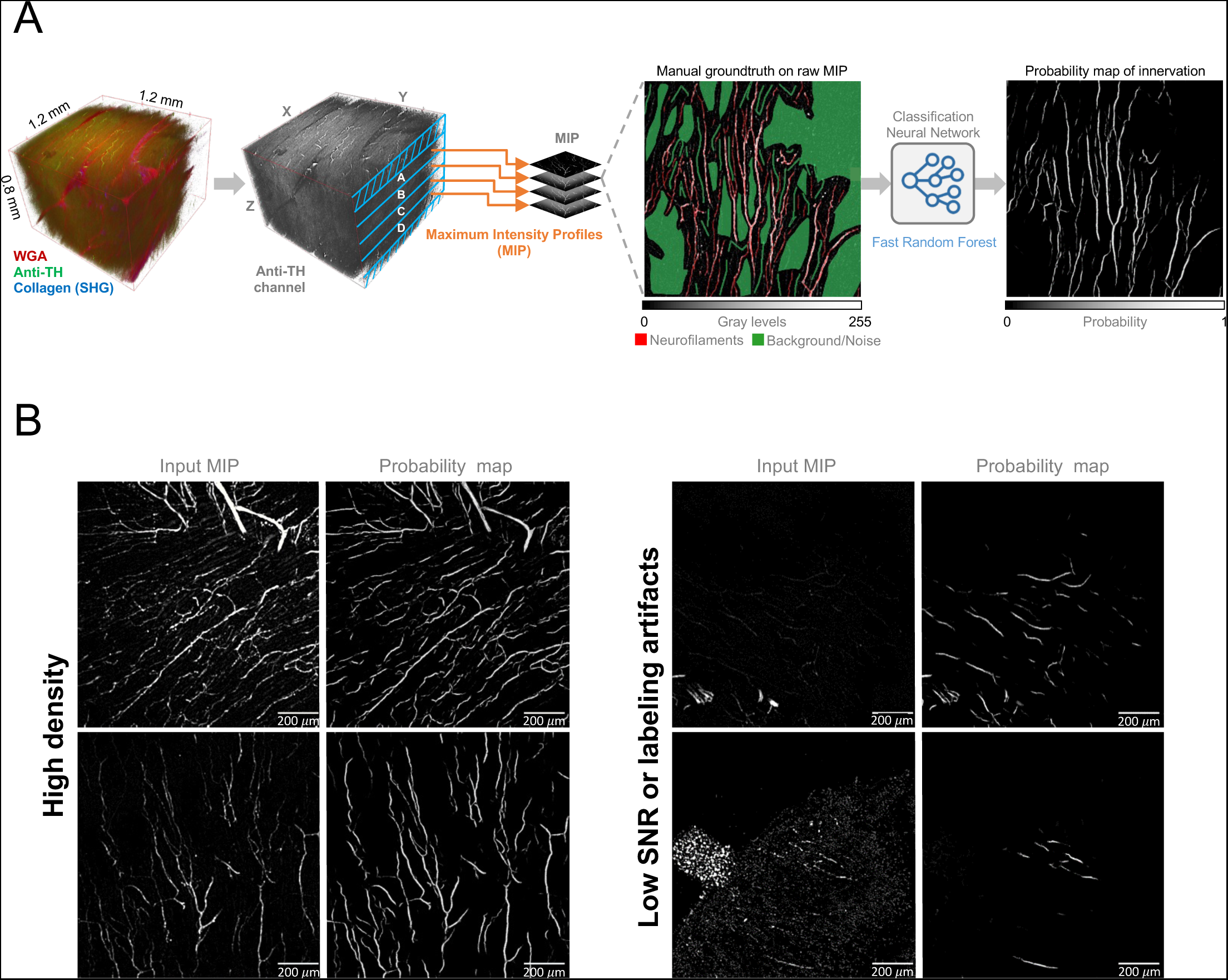
Nerve endings detection software tool. **A)** Schematic representation of the nerve endings detection pipeline. From the left, a representative 3D reconstruction of a three-colors mosaic tile. For each tile, the anti-TH detection channel was isolated, and the 3D volume was projected in 4 maximum intensity projections (MIPs) profiles. Thirty tiles were randomly selected among all the reconstructions and used as a training set: neuronal processes and background were manually classified and exploited to train the software in discriminating the neuronal network and discarding the background noise. Finally, the FastRandomForest method was exploited to generate the nerve endings probability maps. **B)** Examples of nerve endings detection using the developed software tool. The software is able to detect neuronal processes both in tiles with a high density and in tiles with a poor signal-to-noise ratio (SNR), as well as in those presenting artifacts.

**Figure 7:**
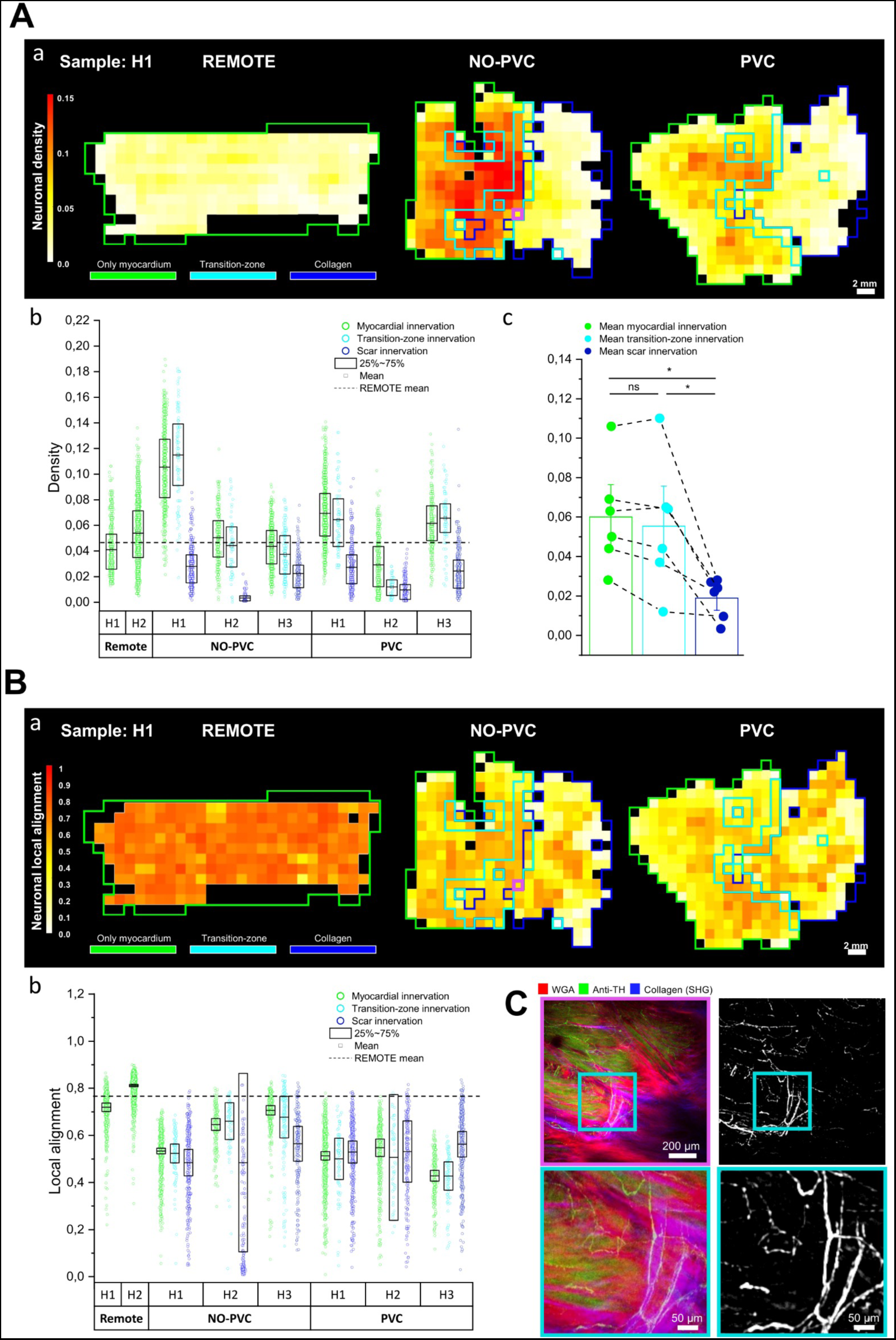
Sympathetic nervous system morphometry. **A)** *Neuronal density*. **Aa)** neuronal density color maps, where myocardium, transition-zone and scar borders have been superimposed, respectively in green, cyan and blue. **Ab)** density values estimated in each tile of every slice corresponding to the myocardial (in green), transition (in cyan) and scar regions (in blue). The dashed line represents the average density value of the two remote sites. **Ac)** mean density values of the corresponding myocardial, transition and scar regions in NO-PVC and PVC slices. One-way ANOVA repeated measures analysis performed with Bonferroni post-hoc testing identified a p= 1 (ns) between myocardial and transition-zone regions, a p= 0.004 (*) between transition-zone and scar regions, and a p= 0.002 (*) between myocardial and scar regions. **B)** *Local alignment of nerve endings*. **Ba)** neuronal local alignment color map, where myocardium, transition-zone and scar borders have been superimposed, respectively in green, cyan and blue. **Bb)** local alignment of nerve endings estimated in each tile corresponding to the myocardial, transition-zone and scar regions. Average values were estimated by weighting the mean by the local density. The dashed line represents the average local alignment value of the two remote sites. **C)** Representative tile of the infarction transition-zone (on the left) and corresponding probability map of the nerve endings (on the right). The corresponding tile is represented in pink in the density and local alignment color maps.

While the extent of neuronal innervation is uniform in the remote site (Figure 7Aa), a higher variability in its density is evident within BZ samples, both in NO-PVC and in PVC sites. First, a low density of nerve endings is apparent in the scar regions with respect to the myocardial regions. In addition, the density of sympathetic innervation in the myocardial areas immediately proximal to the scar tissue is non-uniform with localized high density.

Based on this observation, we quantified neuronal density in the myocardial and scar area of each BZ sample, as well as in the 1mm transition zone between scar and myocardial tissue (Figure 7Ab). Compared to the two remote myocardial sites, the density of neuronal innervation was consistently lower in scar tissue, although nerve endings were not entirely absent (see also Figure 5C), However, we found high variability between the BZ myocardial sites with no consistent differences between PVC and NO-PVC sites. Analysis of neuronal staining within BZ samples (Figure 7Ac) confirmed a significant reduction (68% ± 15%) in the density of sympathetic nerve endings in the scar with respect to the myocardial regions. The transition zone is comparable to the myocardial region as whole, indicating the sharp delineation between scar and myocardial tissue.

Similarly, the local alignment of neurons is uniform in the entire volume of the remote sample (Figure 7B), whereas higher heterogeneity is found in BZ samples, even though no specific differences in local distribution between myocardium and scar can be perceived.

In addition to this higher variability, in BZ samples we found a decreased alignment of sympathetic nerve endings compared to remote sites, both in PVC and NO-PVC sites, and both in the myocardial, transition-zone and scar regions.

## DISCUSSION

### 3D print-guided ex vivo sampling accurately recovers target sites

We report a novel pipeline to perform precise cardiac tissue sampling, of *in vivo* defined sites, which would otherwise not be identifiable in the isolated heart. As proof of concept, arrhythmia-vulnerable sites within the border-zone after MI were sampled. Advanced *in vivo* imaging and functional testing for site identification, and the connection to a 3D printed cast faithfully reproducing the *in vivo* data for sampling, are at the core of this pipeline.

An improved augmented reality system, fusing cMRI-based models with fluoroscopy, was originally developed to support electrophysiology studies and ablation, by providing anatomical landmarks and details superior to fluoroscopy alone^17^. Similar support systems for electrophysiology have subsequently been developed, including CT instead of MR imaging, and coupled with various providers for electrophysiology interventions^23^ or guidance of intramyocardial injections, using the BZ as reference point^18^.

Here we introduced 3D printing, transferring the *in vivo* information to an individualized 3D cast incorporating the location of sites of interest, to allow precision tissue sampling. The cast served as a restraining cradle for the explanted heart and the uniqueness of the geometry of an individualized cast eliminates misalignment. The correspondence in form during sampling and *in vivo* annotation is crucial for high accuracy, for which proper filling of the heart ex vivo is essential.

By retrieving ablation lesions created at pre-defined targets, the 3D print-guided sampling was shown to be feasible and accurate. For the proof-of-concept, the recovery of samples from the MI BZ was also accurate, as supported by the match between samples and *ex vivo* MRI. The size of the arrhythmia-vulnerable sites was more variable, but remained within a 1×1 cm area dictated by the slit that guided the recovery of the samples. Nevertheless, the application to recover arrhythmia-vulnerable sites illustrates some of the remaining challenges of the methods, i.e. to balance sensitivity versus specificity. To ensure inclusion of the region of interest, the excisions of PVC and NO-PVC sites were widened in the circumferential axis, while maintaining the apex-base predefined dimension of the sampling slits in the 3D cast. However, this may reduce sensitivity by including less relevant tissue within the samples. The 3D-printing methods allow substantial flexibility in designing the slits for recovery and could be further adapted, when studying smaller or larger pathophysiological regions of interest.

### A pipeline for microarchitecture analysis from meso-to microscale

For a proof of concept of targeted sampling, we recovered arrhythmia-vulnerable sites from the MI BZ identified as sites where frequent PVC clustered during adrenergic stimulation. We previously have shown that cardiomyocytes in the BZ have an increased sensitivity to adrenergic stimulation with isoproterenol, which could possibly be due to denervation, given that altered innervation is a known feature of the BZ^10^. Therefore, we hypothesized that the arrhythmogenic nature of certain areas of the infarct BZ could rely on the structural differences that may occur during the remodeling process in areas of frequent PVCs with respect to non-arrhythmogenic sites. Testing this hypothesis required the development of a pipeline for preparing large-scale samples for 3D reconstructions at micron-scale resolution, together with tailored analysis tools that would encompass the mesoscale organization of the tissue down to the microscale of local innervation morphometry.

Previous research studied cardiac sympathetic innervation on entire mouse hearts^13,14,24^ or in small 2D sections of human or pig heart tissue^25,26^. Integrating the information power of the human-resembling pig model with the investigation of centimeters-sized samples at subcellular resolution has come at the cost of several optimizations required in the treatment of the samples, as well as long imaging timings and the handling of large volumes of data.

The data illustrates the complexity of the microarchitecture of the BZ. In this ischemia-reperfusion pig model, we identified islands of cardiomyocytes within the scar tissue as well as nerve endings projecting into the scar, possibly connecting to the myocytes. We also showed that the transition zone between scar and myocardium is less than 1 mm, underscoring the abrupt change in tissue properties that likely contributes to the arrhythmia substrate. However, the predictions for a distinct signature of sympathetic network structural remodeling in PVC sites were not confirmed, despite selecting a group of MI animals with the most profound arrhythmias. Whereas this is only a pilot with small sample size, the implications are that differences in ultrastructure are likely subtle and would require a substantial number of animals for further study. Alternatively, another explanation could be that the differences are not in the tissue ultrastructure but could rely on the functional properties of cardiomyocytes or of the sympathetic network in releasing norepinephrine.

### Perspectives for wider application

Some limitations of the in vivo cardiac imaging and registration process could still be improved. Recent advances in imaging, such as single photon counting CT, hold promise to further reduce registration errors due to superior imaging resolution^27^. Accurate fusion of imaging modalities remains an active field of research and the ongoing development may abbreviate the pipeline.

The pipeline for sampling is not restricted to arrhythmia research but can be applied to recover sites of interest characterized in vivo by other functional markers, such as local contraction abnormalities, localized edema or fibrosis, signs of local inflammation, or recovery of treatment sites. Beyond ultrastructure, the analysis can include functional cellular studies. In a pilot study, we found that viable cardiomyocytes could be isolated from the recovered samples, with preserved membrane potential and action potential configuration (data not shown). Furthermore, 3D print-guided sampling could easily be incorporated in other organs and other fields of medical research to spatially match functional *in vivo* investigation with *ex vivo* tissue, cellular and molecular information.

## ONLINE METHODS

### Study design

Supplemental Figure S1. presents the study design. A first series of experiments is conducted in healthy pigs to develop the methodology for targeted sampling and validate the accuracy of the sampling. A second series of experiments applies the novel targeted sampling methodology to an animal model of myocardial infarction. After identifying arrhythmia vulnerable sites, these and matching control sites are sampled for analysis of microarchitecture.

### Animal model

Ten domestic pigs (Strain TN70, Topigs, Norsvin) of both sexes were used in this study. Treatment was in accordance with international guidelines (European Directive 2010/63/EU) and protocols were approved by the KU Leuven ethical committee (ECD137/2018). All *in vivo* studies were performed under full anesthesia and artificial ventilation. Three healthy pigs were used for validation of the accuracy of sampling. In seven pigs, a myocardial infarction was created by 120 min balloon occlusion of the left anterior descending coronary artery followed by reperfusion, according to previously described protocols^10^. One animal died within 24 h after induction of myocardial infarction.

### Anesthesia during imaging and electrophysiology procedures

All *in vivo* studies were performed under general anesthesia and artificial ventilation at 6-10 ml/kg with 50% oxygen-air mixture. Sedation was with tiletamine/zolazepam 8 mg/kg and xylazine 2.5 mg/kg IM, anesthesia was induced with propofol 3 mg/kg and maintained with 10 mg/kg/h IV combined with remifentanil 0.2-0.4 µg/kg/h IV. Antibiotics (pre-operative cefazolin 22-50 mg/kg, IV and post-operative amoxicillin 15mg/kg or enrofloxacilin 4mg/kg, IM) and anticoagulation (acetylsalicylic acid, 500 mg and periodic heparin 5 000-10 000 IU, IV) were routinely administered.

### Cardiac magnetic resonance image acquisition

Magnetic resonance imaging was performed on a 3 Tesla magnet (Magnetom Prisma^fit^, Siemens Healthineers, Erlangen, Germany). Animals were installed in supine position and ventilated throughout the procedure. In addition to the spine coil 2 phased-array coils were used to cover the entire anterior and bi-lateral aspects of the convex thorax for better signal to noise ratio. All images were acquired with ECG gating where appropriate and under suspended respiration (end-expiratory/open mouth).

Cine images were acquired in 6 long axis (LAX) slices centered and equally distributed (every 30°) around the LV central axis (line connecting the apex end the midpoint of the mitral ring, in addition to a contiguous stack of short axes (SAX) covering the complete LV atrium and ventricle (+/− 20 slices) and planned perpendicular to the former. All cine imaging was performed using a segmented 2D balanced steady state free precession sequence with a slice thickness of 6mm and aiming at an in-plane pixelsize of 1.3×1.3 mm (adapting matrix size and field-of-view (FOV) for the pig), using retrospective gating to reconstruct 30 phases per cardiac cycle. Typical sequence parameters are: echo time: 1.58ms; repetition time: 39.71ms; flip angle: 50°

A bolus of Gd (gadoterate meglumine, Dotarem, 0.2 mmol/kg) was given at a rate of 5 ml/s using a pressure injector to achieve a high concentration at first pass to give sufficient blood-tissue contrast. To capture the ascending-arch- and descending aorta, a modified vertical long axis or 5-chamber view was extended in the sagittal plane. An angiogram was acquired during the first passage of contrast.

Fifteen minutes after contrast injection a 3D inversion-recovery sequence was applied in SAX (+/− 20 slices, covering the same area (LV and LA) as the cine SAX slices) and LAX stacks (8 slices, covering the scar) to define the infarcted myocardium.

### CMRI image processing to 3D anatomical models

The cMRI images were processed offline using custom Matlab software (RightVol, KU Leuven, Leuven, Belgium). The left ventricle (LV) endocardium, the endocardial right side of the septum and the whole heart epicardial surface were manually segmented by contouring the LGE short-axis images (Supplemental Figure S2A), including papillary muscles in the cavity. To aid the orientation and registration of the LV during the in vivo study, the aorta and papillary muscles were integrated in the models. For this purpose, the aorta was automatically segmented from the 3D angiogram (Supplemental Figure S2B) by applying a thresholding algorithm and papillary muscles were automatically segmented within the cavity by a binary segmentation (Otsu) of the SAX LGE images (Supplemental Figure S2C). The contours were then transformed into 3D masked images that were used to create 3D surface models using a binary thresholding algorithm implemented in MeVisLab (Bremen, Germany). (Supplemental Figure S2D-E).

### *In vivo* fluoroscopic imaging and cMRI model integration

The fluoroscopic setup consisted of 2 separate fluoroscopes positioned in an 80° biplane. The right anterior-oblique (RAO) fluoroscope (mobile OCE Fluorostar, GE Healthcare, Germany) was positioned at 30°, and the left anterior-oblique (LAO) fluoroscope (Coroskop HiCor, Siemens Healthineers,Erlangen, Germany) was positioned at 50°with 15° caudal angulation.

The custom software application (LARCA) was used to integrate the 3D anatomic models into the biplane fluoroscopy framework for live augmented reality imaging and catheter guidance^17,28^. To correctly integrate the preoperative CMR-derived 3D models into the intraoperative fluoroscopic setup, we calibrated the imaging parameters of both fluoroscopes similar to previously described^28^. Non-linear distortions were determined by imaging a square wire grid with 1-cm pacing. The crossing points of the grid visible in both views was used to obtain pairs of distorted-undistorted coordinates. The distorted-undistorted coordinates allowed the determination of the distortion error by back-projection to an ideal square grid projection^17,28^. Calibration of the weak perspective model^28^ and registration between the 3D cMRI-derived LV surface model and the ventriculogram was performed similar to ^28^ and ^18^, respectively. The calibration was further validated using a 3D printed phantom of the LV endocardial cavity of a control pig on which 9 iron beads (diameter 1mm) were placed around the LV. A CT scan and reconstruction of this phantom was used as the gold-standard and several fluoroscopic images of the phantom with indicating the beads in different poses were used to determine the translation and angulation error (Figure S4).

The 3D models were rendered transparent and were superimposed on the fluoroscopic images. The models could also be manually adjusted to minimize the registration error and offset as determined by the operator. The position of the 3D models was then fixed and live overlay was applied to fluoroscopic views.

### Creating endocardial RF ablation lesions

Under general anesthesia, the femoral vein, jugular vein, and carotid artery were cannulated with 7-8Fr sheaths. Standard EP catheters (Decanav, Biosense Webster, Irvine CA, USA) were positioned in the RV and coronary sinus. To approximate the cMRI cardiac dimensions, pacing was performed from the right atrium or RV at the cMRI heart rate. A bi-plane left ventriculogram was obtained with 70-110mL 50-80% contract-saline mixture delivered via a Medrad pressure injector (Bayer, UK) and pigtail in the LV apex, and end-diastolic images were imported using a framegrabber. The multimodal imaging allowed guidance of the catheter to the predefined target sites. Ablation was achieved by an RF generator (Stockert, Biosense Webster, Irvine CA, USA), with settings for irrigated tip ablation of 30W, 45-50°C and irrigation at 17mL/min. If VF occurred, the animal was defibrillated with a 200J biphasic shock and the 3D model fusion redone.

### Electroanatomical mapping under adrenergic stimulation

An EnSite Precision 2.0 mapping system (Abbott Medical, New Jersey, USA) paired with a BARD Electrophysiology Labsystem Pro EP (Boston Scientific, USA) was used for electroanatomical mapping as described previously^10^. The EnSite multi-electrode array was positioned centrally in the LV via arterial access and standard EP catheters in the RV and coronary sinus via venous access. A detailed elecroanatomical map was created by roving the ventricle and annotating the peak-to-peak electrogram voltage measured using a mapping catheter (Lasso or Navistar, Biosense Webster, Belgium).

Animals underwent electrophysiological testing by performing adrenergic stimulation using isoproterenol infusion. First, a baseline recording was made without isoproterenol infusion. Thereafter, isoproterenol was infused in 4 incremental doses of 0.01 µg/kg until reaching 0.04 µg/kg, each dose over 5 minutes. During the 20-min isoproterenol infusion protocol a continuous non-contact mapping recording was made. MI animals exhibited an isoproterenol dose-dependent increase in heart rate with an increase in PVC frequency (consistent with previous work^10^).

### Construction of 3D-print model for targeted sampling

A unique cast model for 3D printing was constructed based on a precision 3D modelling workflow in the open-source Blender software (v2.9.0). The workflow started from the anatomical 3D surface models of the epicardium (Figure S2E). The epicardial mesh was outwardly extruded for 4 mm to create a cast of the heart. Then the intersection of the cast with a plane positioned at the intra-ventricular groove was calculated (referred to as dividing plane). This intersection was used to divide the cast in a left and right ventricular part. Thereafter, handles were added to the left and right ventricular part of the cast. This was done by creating the intersection of a plane parallel to the dividing plane on each side (referred to as left and right handle plane resp.). Four nodes were added to each plane and were connected to create the handles (to be precise, 2 × 4 nodes in the dividing plane). Thereafter the septal plate was created. To create the septal plate we restarted from the epicardial mesh and joined the segmentation of the right ventricular septum (Figure S2E) to create a combined mesh. The intersection of the combined mesh with the left-hand plane and the dividing plane was determined. We hid the intersecting vertices which resulted in disconnected regions. For the epicardial mesh fragment of the combined mesh this results in three disconnected sub-meshes. The portions that were not between the 2 planes were removed. For the right ventricular septal mesh part of the combined mesh, we removed the region that was between the 2 planes. The last step consisted of creating the planes of the septal plate.

The slits for targeted sampling were created by performing a Boolean operation between a beam centered at the sampling target and the above created cast. In each sampling a beam was created with a cross-section of 1×1 cm and a length of 5 cm. The center of mass of each beam corresponded with the targeted sampling point. The beam was kept parallel to the XY-plane which corresponds to the short axis image plane determined during in vivo cMRI. The beam was rotated around the Z-axis so that the longest was perpendicular to the endocardial surface.

### 3D printing of casts for sampling

The 3D cast with slits for targeted sampling was constructed from the cMRI epicardial surface model using precision 3D modelling in the Blender software (v2.9.0). The casts were printed using a commercial 3D printer (Builder Premium Large, Builder, The Netherlands), with a 0.4mm nozzle size in standard PLA material (PLA, Builder, The Netherlands) and using the following settings: layer height of 0.2mm, with a print bed temperature of 60°C and a nozzle temperature of 210°C.

### Sacrifice and preparation of the heart for sampling

For sacrifice, animals were sedated and anesthesia was induced with pentobarbital (20 mg/kg pentobarbital, IV) under artificial ventilation. Through a medial sternotomy, the ascending aorta was cannulated with a cardioplegia needle and sheath, and heparin (5 000 IU) administered and the aorta cross clamped. The superior- and inferior vena cava were also cross clamped to achieve only coronary-pulmonary circulation and volume was vented through an incision in the right atrial appendix. 1L of cold cardioplegia was then infused antegrade to arrest the heart in relaxed state. The heart was quickly harvested, including the great vessels, and washed in cold cardioplegic solution. The great vessels were sutured, and the heart refilled to volumes matching the cMRI used to create the models and 3D prints and sealed to maintain these volumes.

### Ex vivo cMRI

Animals with MI underwent additional *ex vivo c*MRI imaging. Therefore, the MI animals received a 0.2mmol/Kg bolus Gd 15 min before harvesting the heart, which was then prepared as above and submerged in a plastic container filled with oxygenated cardioplegia. This container was then inserted in a phased array knee coil in the 3T scanner and surrounded with cold packs to maintain the temperature below 5°C. The cMRI images were acquired using T1-weighted gradient echo sequence with a resolution of 0.48×0.48×0.50 mm^3^.

### Targeted site sampling

Sampling in the control animals took place directly after refilling the heart. Sampling in the MI animals took place after *ex vivo* cMRI. The marked sample sites were excised and fixed in 4% paraformaldehyde for 8 h, and subsequently stored in phosphate buffered saline (PBS). Samples were cut using a vibratome in the apex to base direction from endocardial to epicardial in 500 µm thick slices.

### CLARITY protocol

The selected slices were washed in PBS 0.1x at room temperature (RT) for 24 h in gentle shaking. Then, the specimens were mounted in a custom-made chamber device using two cover glasses (UQG optics, 60×60 mm, UV fused silica), custom-made 500-μm-thick frame spacers and a bi-component glue (Twinsil speed, Picodent GmbH, Germany). Each sample was placed on a cover glass together with a spacer, carefully covered with a second glass to avoid the formation of bubbles and then the device was sealed with the bi-component glue. It was then filled with 1.5-2 mL of hydrogel solution (4% Acrylamide, 0.05% Bis-acrylamide, 0.25% Initiator AV-044 in 0.01 M PBS) injected from the spacer hole using a 1.0 mL syringe and a 0.45 mm needle. The device was placed in a vial, filled with Hydrogel solution to be completely covered. The sample was incubated for 3 days at 4°C in shaking. After 3 days, the samples were de-gassed using a drier (KNF Neuberger, N86KT.18) and the oxygen was replaced with nitrogen. To aid gel polymerization, the samples were kept at 37° C for 3 h. Finally, they were carefully removed from the sandwich divide and incubated in 30 mL of CLARITY clearing solution (200 mM Boric acid, 4% Sodium Dodecyl-Sulfate (SDS) in deionized water; pH 8.6) at 37° C with shaking to complete clearing of the tissue.

### Immunostaining and samples mounting

CLARITY-cleared slices were washed in RT PBS for 24 h during continuous shaking and then in room temperature PBS + 0.1% of Triton-X (PBS-T 0.1x) for 24 h during continuous shaking. They were then incubated in 1:200 anti-tyrosine hydroxylase (anti-TH; Millipore, Billerica, MA, USA, AB152) in PBS-T 0.1x at 4°C with shaking for 3 days. The samples were then washed in PBS-T 0.1x for 24 h at RT with shaking and, the following day, incubated in 1:200 anti-rabbit Alexa Fluor 488 (Goat anti-rabbit, AB150077; Abcam) and 1:100 Wheat Germ Agglutinin (WGA)-Alexa Fluor 594 (Thermo Fisher, W11262) in PBS-T 0.1x for 2 days at RT in shaking. The specimens were washed in PBS-T 0.1x for 24 h at RT with shaking and, the following day, fixed in PFA 4% in PBS for 15 min and washed in PBS for 5 min three times. Finally, they were incubated in 5 mL of EasyIndex (Life Canvas Technologies) for 2 h to favor the homogenization of the refractive index (RI). The same chamber approach described above was used to mount the samples for the imaging, but, in order to match the RI of the cleared-samples incubated in EasyIndex solution (RI= 1.465), a 500 μm-thick quartz glass (60×60 mm) was used to replace the cover glass on the top. The device was then secured and held stable in a dish filled with a solution of 91% of glycerol in ddH2O (RI =1.465).

### Two-photon fluorescence and second-harmonic imaging

For the image acquisition of the cleared slices we used a custom-made non-linear microscope), as previously described by Olianti* et al.^29^. The laser was focused on the specimens with a tunable refractive index 10× objective (Nikon, CFI 10X G, Plan Apo A.N.0,5 WD 5,5mm, R.I. 1,33-1,51). The described system allowed the reconstruction of the entire surface and thickness of the samples by acquiring serial tiles of 1.2 × 1.2 mm at 512 × 512 pixel (pixel size of ≍ 2.34 μm) and with a Z-step of 5 μm. The serial Z-stacks of adjacent regions were acquired with a lateral overlap of 120 μm. Finally, the raw images were subjected to vignetting correction, performed using the retrospective CIDRE (Corrected Intensity Distributions using Regularized Energy minimization) algorithm^30^ and stitched together using the Fusion plugin of the commercial Huygens software 21.10 (Scientific Volume Imaging, The Netherlands).

### Estimation of the CLARITY-related samples swelling in XY

In each sample, we evaluated the CLARITY-related tissue swelling in the two dimensions (X,Y) both in the myocardial region and in the scar region. To these ends, we photographed the samples before and after the clearing procedure on a 1-mm graph paper sheet. For each sample, we estimated the expansion in the two axes by measuring the length of the myocardial and the scar regions in several positions of the sample, before and after the clearing procedure. The corresponding X and Y expansion coefficients were applied to resize the mosaic reconstructions to their original dimensions before performing further analyses.

### Segmentation of myocardium, scar and transition-zone

We segmented myocardial and fibrotic tissue in the slices involving muscle autofluorescence and collagen detection channels. Frame by frame, we down-sampled images to a pixel size of 280 × 280 µm and we applied a threshold-based segmentation using the AutoThreshold plugin of Fiji software^31^. We thus created 3 masks corresponding to myocardial, scar and overlapping areas (defined as the overlapping area between myocardium and scar segmentations). We calculated the volume of myocardium, fibrosis, and overlapping area by multiplying the number of voxels for the volume of each voxel (0.000392 mm^3^). Finally, volume of myocardium, scar, and overlapping areas is re-scaled accordingly with the expansion factor previously estimated. The percentage of collagen is finally calculated for each reconstruction as follow:

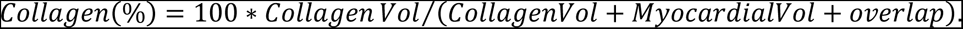

We also identified the transition-zone by down-sampling the segmentations to a pixel size of 1.2 × 1.2 mm (the dimension of a tile in the XY plane). We merged the segmentations of scar tissue and overlapping areas, thus identifying all the scar tissue, and we expanded them by a “dilate” morphological operator with kernel = 1 pixel. A final logical operator “AND” is applied between the resulting mask and the segmentation of the myocardium, to isolate tiles defined as transition-zone. Additionally, we quantified the irregularity of the interface between the fibrotic tissue and the myocardium by measuring the surface area of the scar-myocardium interface as a percentage of the total slice volume.

### Analysis of the sympathetic innervation

Immunostaining for tyrosine hydroxylase identified the nerve endings of the sympathetic system within the heart. Sympathetic innervation was analyzed in terms of density of nerve endings and local alignment of their orientations. In detail, we developed an automatic pipeline to detect sympathetic fibers in the high-resolution tiles, measure local density and coherency (defined as the degree of local co-alignment) and to reconstruct the 3D map of these structural parameters of the entire slices. In order to better take account of the separation between myocardium and scar regions, we divided each mosaic reconstruction into four different z-sections.

Figure 6A summarizes the automatic detection software of nerve endings. For each tile, neuronal detection channel was selected, filtered to remove noise and background contribution, and split in layers of 150 μm of thickness, thus generating four sub-volumes. For each sub-volume, we squeezed the signal by means of a maximum Intensity profile (MIP) along the z-axis and we identified neuronal filaments through an optimized 2D segmentation procedure based on a neural network, involving the Trainable Weka Segmentation plugin available in Fiji. Thirty tiles were randomly selected among all the reconstructions and used as a training set, where nerve endings and background were manually classified to train the software in discriminating the neuronal network and discarding the background noise. The model assigns to each pixel of the MIP a probability *p* ∈ [0, 1] to be classified as neurofilaments, generating a reliable map of nerves of each subvolume of each tile. Innervation density was calculated by averaging the probability value in each tile. We then exploited the estimated expansion coefficients of each sample to scale the results accordingly. Neuronal local alignment was calculated by applying the plugin “OrientationJ Measure” of Fiji^32^.

The resulting density and local alignment values were remapped into matrices, where each element spatially correlated to the corresponding tile of the mosaic. We then used the myocardium, scar, and transition-zone segmentations to mask the matrices and discriminate the values corresponding to the three regions of interest. To balance the impact of the actual amount of innervation in the local alignment estimation, we calculated the average value of density-weighted neuronal alignment and its density-weighted standard deviation.

### Analysis and statistics

Off-line calibration and data analysis was performed with the operator blinded. All data are presented as mean ± SD. Data were compared with Student’s t-test, or ANOVA with Bonferroni post-hoc testing, as applicable and indicated in the figure legends.

## ACKNOWLEDGEMENTS

We would like to thank Patricia Holemans and Roxane Menten for the assistance during experimental procedures.

PC, RW, LR and PC disclose support for the research of this work from FWO, the Fund for Scientific Research-Flanders [grant number G097021N] and PC and RW disclose support from KU Leuven BOF-C1 [grant number C14/18/079].

## AUTHOR CONTRIBUTIONS

DV, MA, RW, KS and PC were responsible for conceptualizing and DV, MA and PC performed the targeted sampling experiments. CO, FG and LS conceived and designed the structural study of sampled tissues. CO was responsible for optimizing and performing the clearing, staining, mounting and fluorescence imaging of the sampled tissues, as well as performing the structural analyses. FG was responsible for developing the software tools and pipelines for structural analyses. SDB was involved in setup and tuning of the LARCA workflow as well as the printing of the PLA casts. CKN and LR were involved in antibody optimization and validation, tissue sectioning and sample selection. DV, CO, FG and PC prepared the figures. KS, LS, RW and PC contributed to the essential material, models and facilities. DV, CO, MA, FG, LR, KS, LS and PC were involved in preparation of initial the manuscript. All authors, except MA, were involved in final revision of the manuscript.

## Supplemental Data

**Figure S1.**
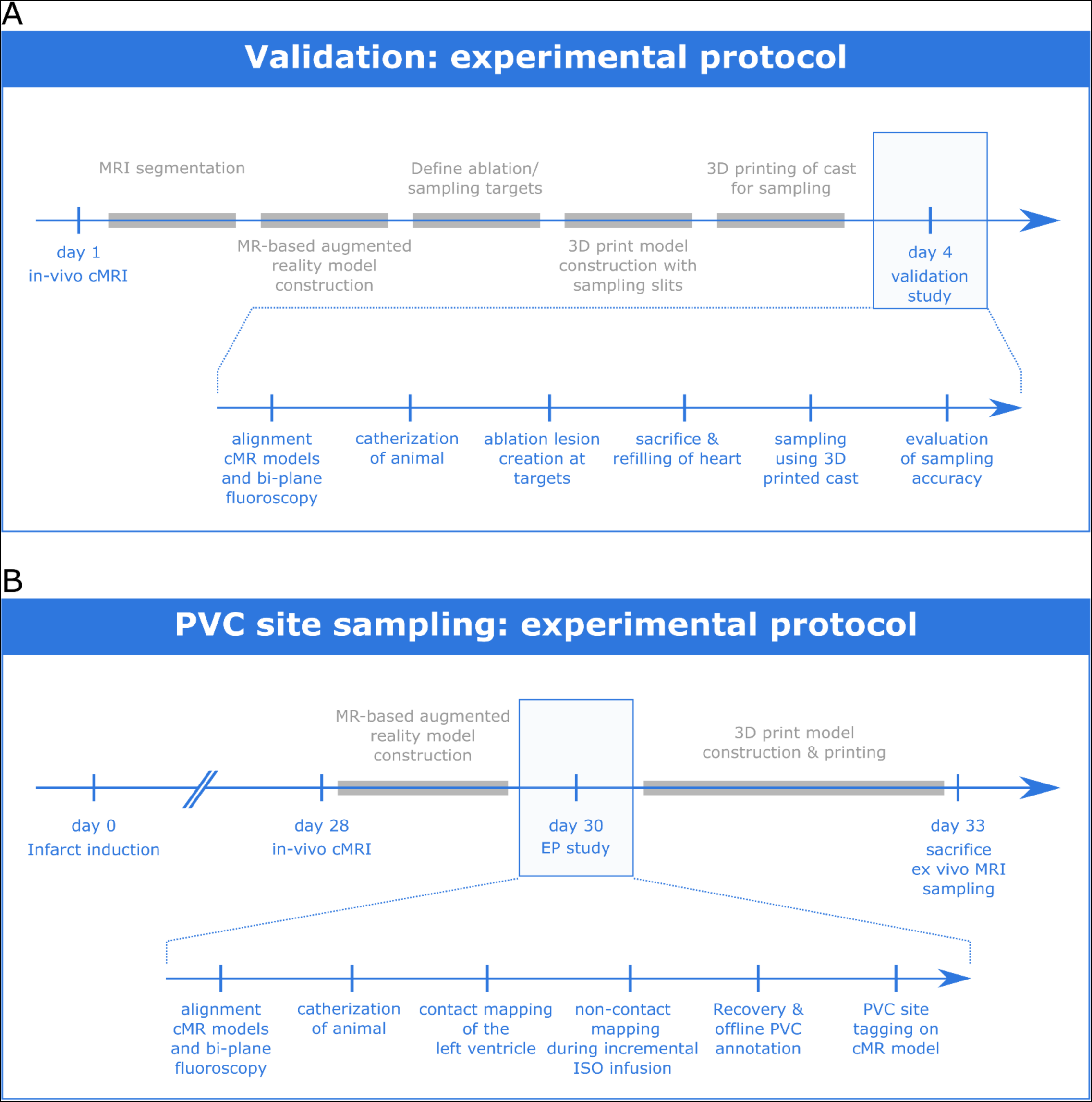
The experimental protocols for the targeted sampling validation and arrhythmia-vulnerable substrate sampling studies. **A)** The experimental protocol for the validation of the targeted sampling methodology. The blue markers indicate experimental procedures, the grey boxes indicate offline analysis performed between experimental procedures. **B)** The experimental protocol for sampling of in vivo identified vulnerable arrhythmia substrate. Color code is similar to panel A.

**Figure S2.**
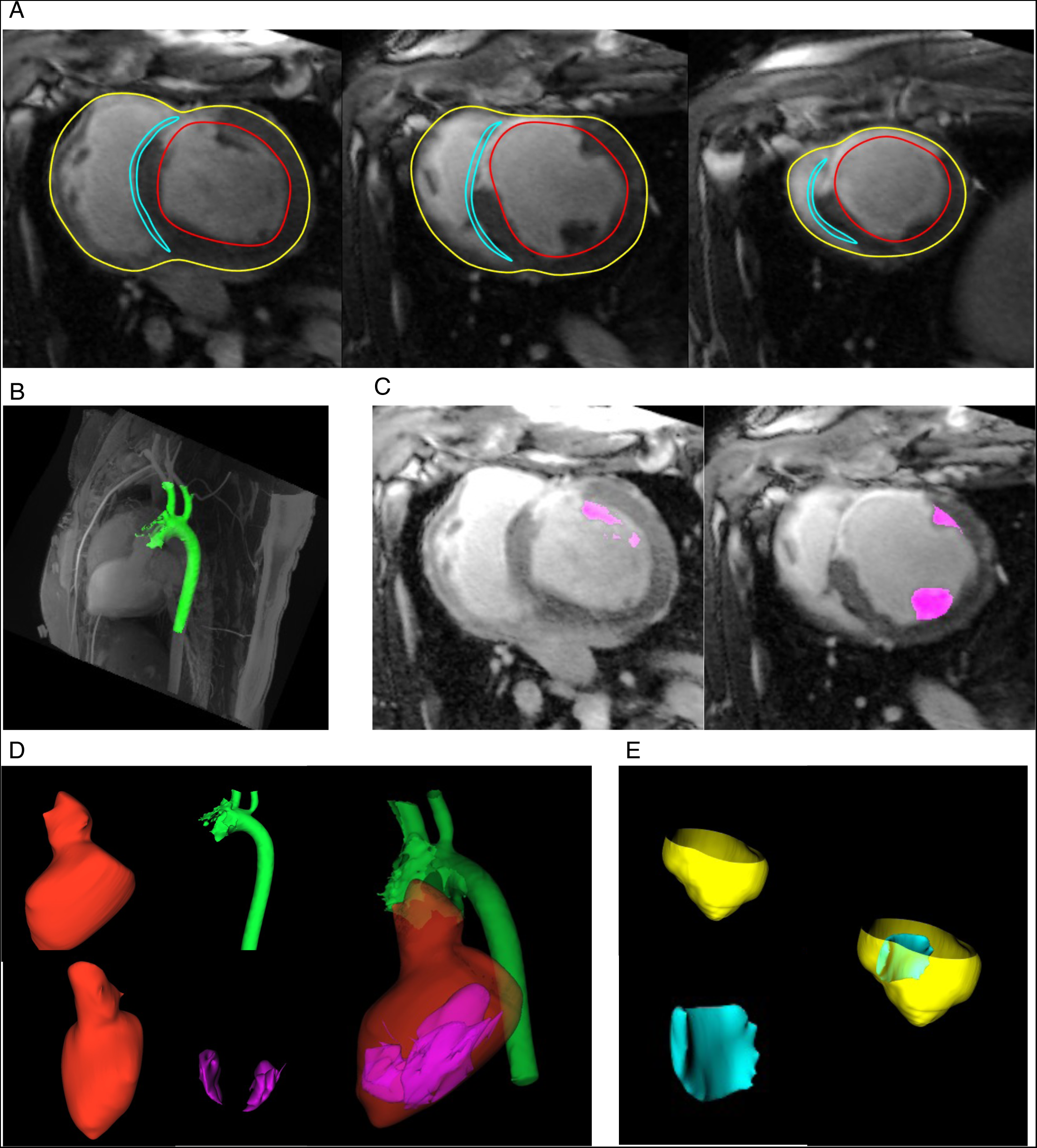
Cardiac magnetic resonance image segmentation and 3D model construction. **A**) Illustration of representative basal to apical short axis LGE slices and superimposed manual segmentation of the LV endocardium (red), epicardial surface (yellow) and RV septum (cyan). **B**) Maximum intensity projection of the 3D angiogram with aorta model obtained by automated thresholding. **C**) Papillary muscle segmentation (magenta) by thresholding within the cavity after endocardial mask from the LGE short axis images. **D**) Resulting models for AR system. Left, the LV endocardium. Middle: corresponding model of the aorta (top) and papillary muscles (bottom). Right: final integrated models. **E**) Resulting models for CAD to produce cast. Left: epicardial model (yellow, top) and RV septal model (cyan, bottom) Right: integrated models.

**Figure S3.**
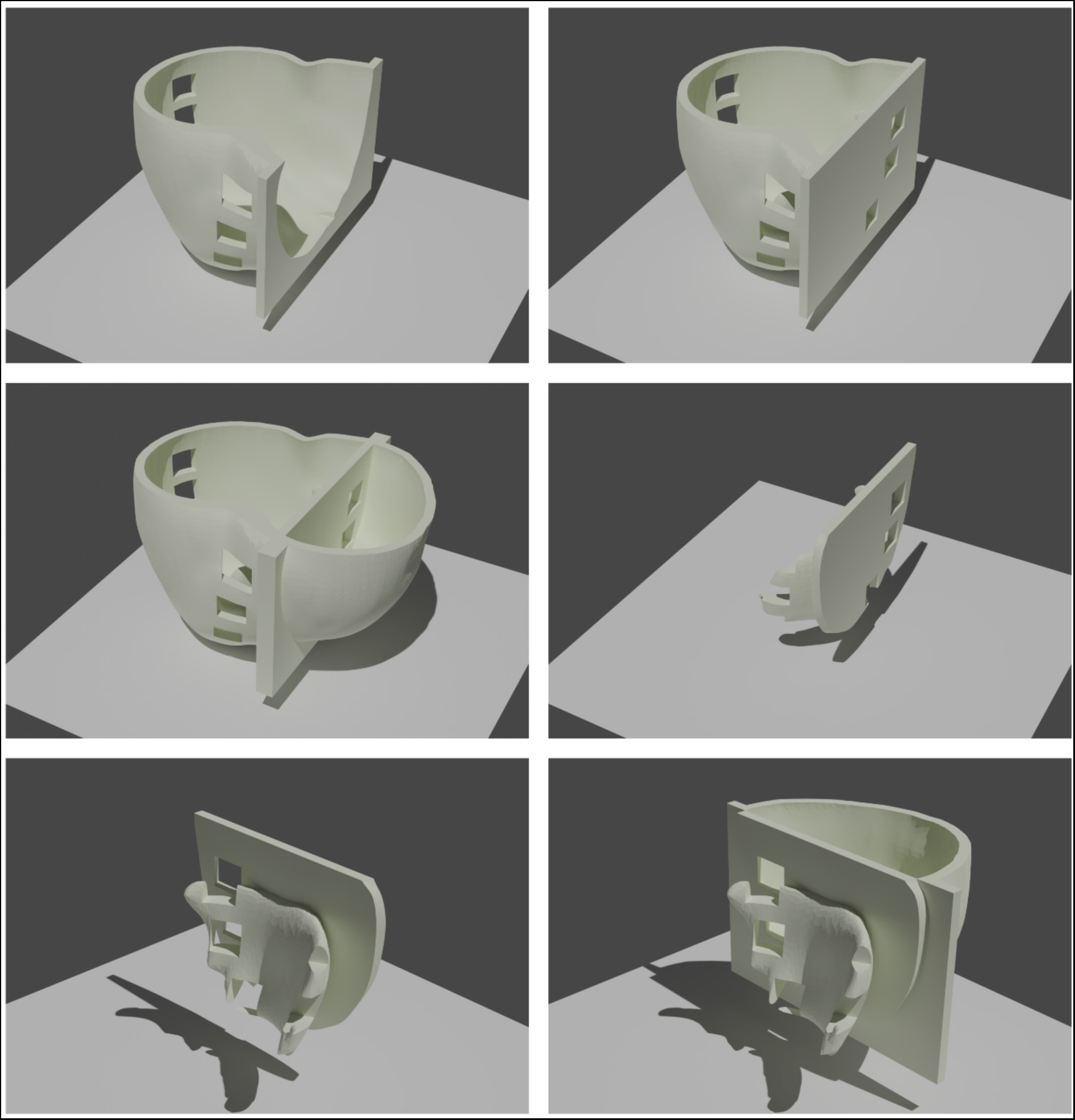
Rendering of the 3D printing design for targeted sampling. Different views and compositions of the three components (left ventricle cast, right ventricle cast and septal plate) making up the design for targeted sampling.

**Figure S4.**
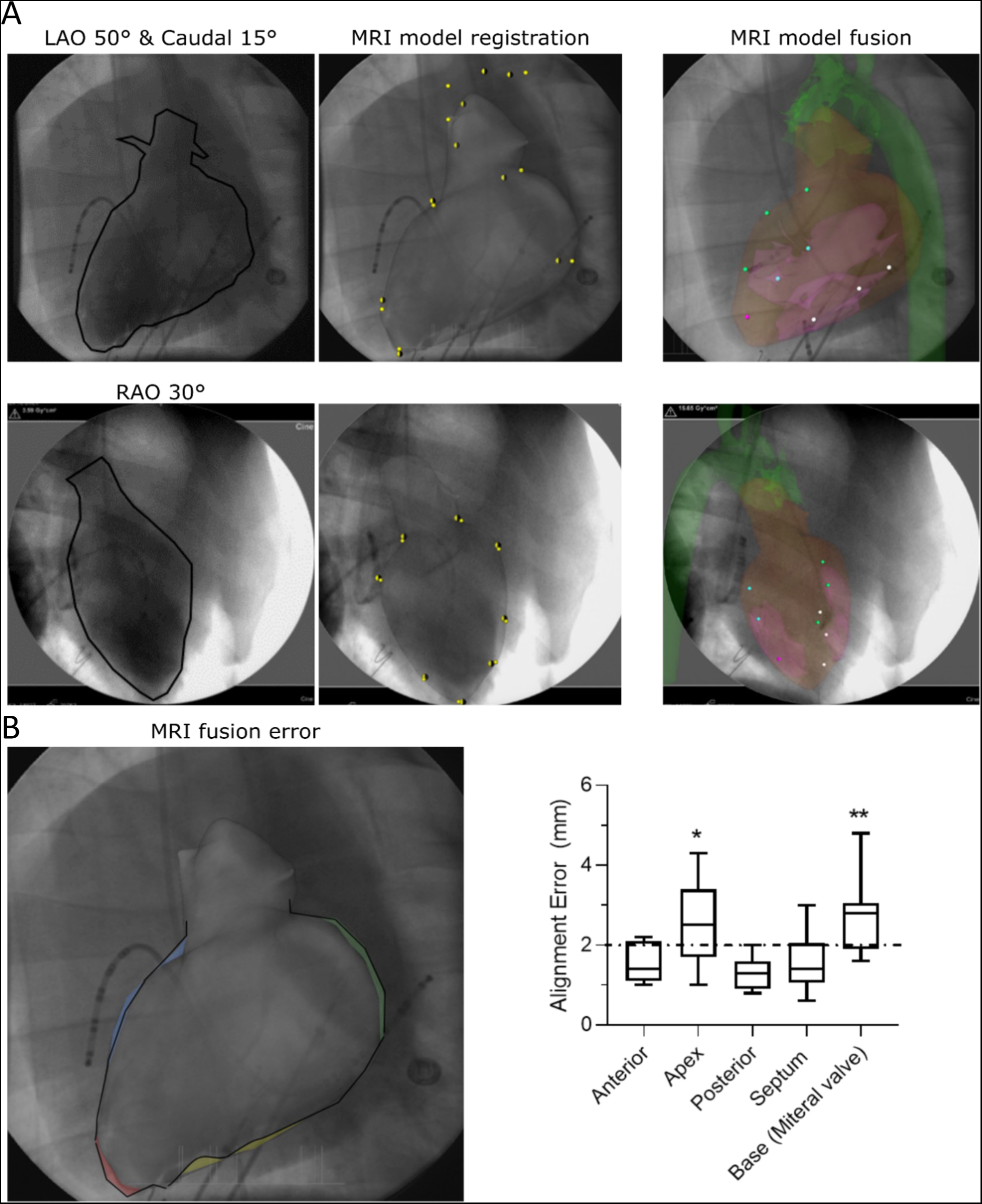
Augmented reality angiography (LARCA) and 3D model catheter guidance. **A**) Co-registration of the magnetic resonance imaging (MRI) based anatomical models with biplane fluoroscopy was performed using an iterative closest point (yellow vs. black dots) cloud registration. Top and bottom show the 2 different fluoroscopes. The panel on the left shows the fluoroscopic contrast based contour. The middle panel shows the points (yellow) used for co-registration of the MRI model. The right panel shows the augmented fluoroscopic view. **B**) Representative example of assessment of the alignment for LAO view (left) and mean data for all views (N_pigs_=9, right).

**Figure S5:**
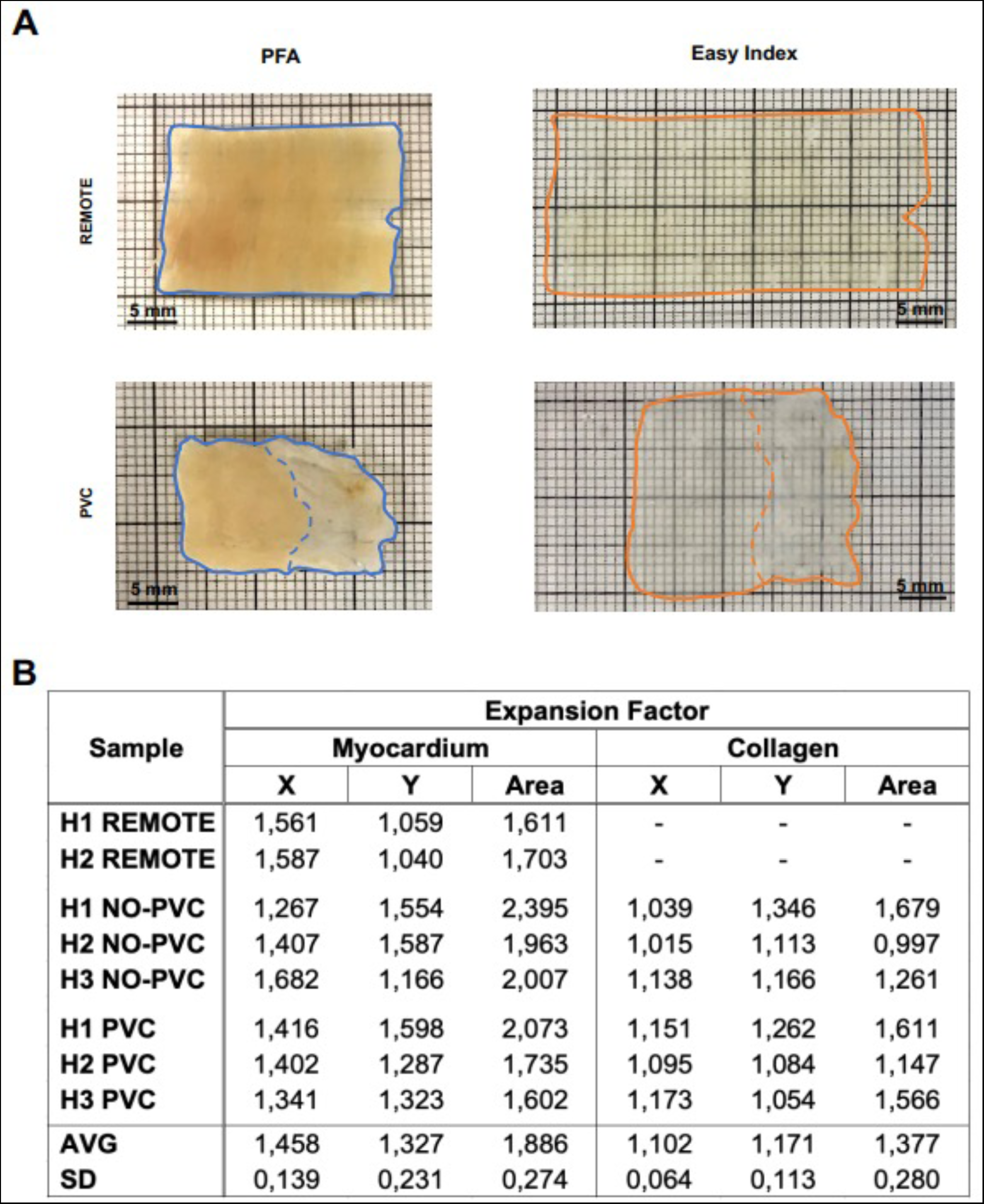
Estimation of the CLARITY-related expansion coefficients. **A)** A representative remote and PVC site, both before and after the clearing protocol. The outer borders of the slices have been highlighted in blue (before the clearing) and orange (after the clearing) whereas the dashed line highlights the border between the myocardial and scar regions. **B)** Expansion coefficients of the myocardial and scar regions calculated for each sample in the XY axes and in the surface area. Last rows indicated the average (AVG) and standard deviation (SD) over all samples.

**Supplemental video V1:**
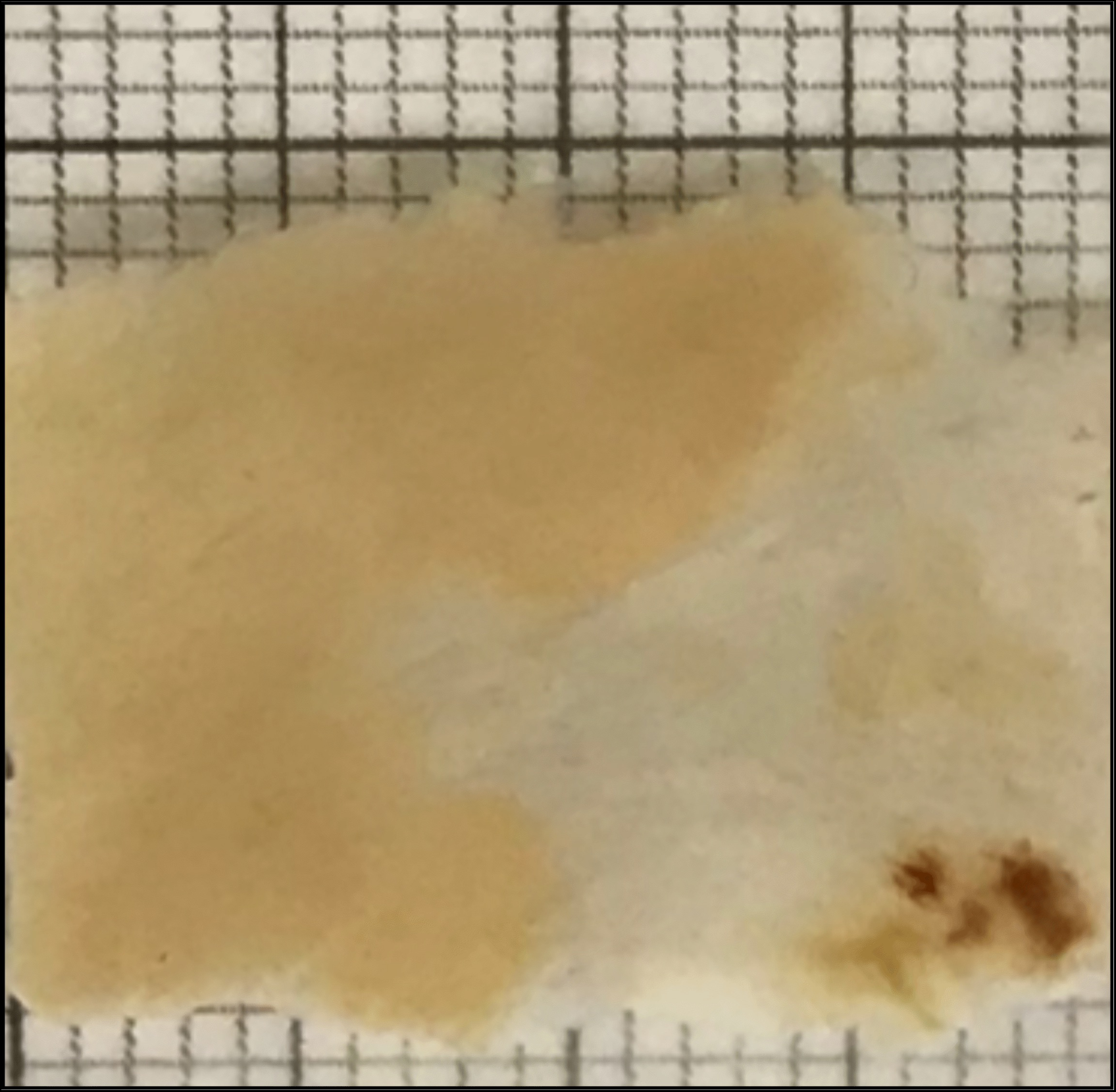
Overlay between targeted sample and high-resolution ex vivo MR image from matched region.

**SUPPLEMENTARY TABLE T1:** CMR DATA ON STRUCTURE AND FUNCTION IN CONTROL PIGS AND 1-MONTH AFTER MI.

|  | MI (N<sub>pigs</sub> = 6) | Control (N<sub>pigs</sub> = 3) | P value |
| --- | --- | --- | --- |
| Infarct size (%) | 18.8 ± 3.5 | 0 ± 0 | <0.0001 |
| Ejection fraction (%) | 40.4 ± 9.6 | 68.0 ± 3.9 | <0.01 |
| End-diastolic volume (mL) | 148.7 ± .0 | 82.7 ± 40.6 | <0.05 |
| End-systolic volume (mL) | 117.1 ± 27.3 | 40.3 ± 13.1 | <0.05 |
| Left ventricular mass (g) | 147.4 ± 12.8 | 131.7 ± 18.6 | 0.7674 |
| Heart rate (beats per minute) | 88.8 ± 12.2 | 82.7 ± 28.2 | 0.8825 |

